# A computational neural model for mapping degenerate neural architectures

**DOI:** 10.1101/2020.11.13.382192

**Authors:** Zulqarnain Khan, Yiyu Wang, Eli Z. Sennesh, Jennifer Dy, Sarah Ostadabbas, Jan-Willem van de Meent, J. Benjamin Hutchinson, Ajay B. Satpute

## Abstract

Degeneracy in biological systems refers to a many-to-one mapping between physical structures and their functional (including psychological) outcomes. Despite the ubiquity of the phenomenon, traditional analytical tools for modeling degeneracy in neuroscience are extremely limited. In this study, we generated synthetic datasets to describe three situations of degeneracy in fMRI data to demonstrate the limitations of the current univariate approach. We describe a novel computational approach for the analysis referred to as neural topographic factor analysis (NTFA). NTFA is designed to capture variations in neural activity across task conditions and participants. The advantage of this discovery-oriented approach is to reveal whether and how experimental trials and participants cluster into task conditions and participant groups. We applied NTFA on simulated data, revealing the appropriate degeneracy assumption in all three situations and demonstrating NTFA’s utility in uncovering degeneracy. Lastly, we discussed the importance of testing degeneracy in fMRI and the implications of applying NTFA to do so.

## 1 Introduction

Degeneracy refers to the capability of different structures to produce the same effects [1, 2, 3]. For example, different sets of codons in genetics can produce the same phenotype [4]. Different ion channels - more than are strictly necessary - are used to tune the firing rate of neurons [5]. Different distributions of neural modulators and circuit parameters nonetheless produce the same rhythmic activity in a neural circuit [6, 7]. Simple motor behaviors, like finger tapping, may also be produced by an abundance of distinct motor pathways [8, 9, 10, 11]. In functional neuroanatomy, degeneracy refers to the notion that the brain may have multiple solutions or a surplus of neural pathways to produce the same mental state or behavior [12, 13, 14]. Indeed, computational simulations show that degeneracy is high in networks with high complexity such as the brain [15, 16], in which multiple distinct, parallel structural pathways may lead from a source node to a destination node. Such an architecture enables a degree of robustness to changes in the neural environment (e.g. due to tissue damage) [12, 14]. ^1^

In cognitive neuroscience, degeneracy in functional neuroanatomy suggests there might be systematic sources of variance across trials or individuals that are of interest for the brain-behavior relationship. For example, two individuals may use different neural pathways to perform the same task, or one individual may use different neural pathways in different moments when performing a task. Commonly used analytical approaches often treat such variation across trials within a condition and across individuals within a sampled group of participants as error. For example, functional neuroimaging studies that examine task-dependent changes in functional activation often estimate parameters assuming invariance across trials or participants. Offering a bit more flexibility, recent machine learning approaches have also been applied to functional neuroimaging data (e.g. multivoxel pattern analysis)[18, 19], however, these approaches commonly rely on supervised analytical approaches that imply a common neural activation pattern for trials in the same task [20]. In both cases, summaries are calculated either across participants, trials, or both in order to increase signal-to-noise ratios, and residual variance is assumed to provide an estimate of error for calculating inferential statistics. However, in doing so, these approaches are assuming a non-degenerate functional architecture *a priori*. As a result, little is known about the extent to which these assumptions prevail vs. the extent to which there is degeneracy in functional neuroanatomy.

Uncovering degeneracy requires analytical tools that are explicitly designed for this purpose. If the brain provides multiple solutions to complete a given task, then functional activation patterns in a given study may depend on the participant and moment in time (i.e. by stimulus or trial) in ways that are unbeknownst to investigators. Thus, it is important to develop an analytical approach that can identify sources of structure in signal with minimal supervision - that is, without relying on strong *a priori* assumptions of investigators of how functional activity ought to relate with task performance. Here, we propose a novel computational model, referred to as Neural Topographic Factor Analysis (NTFA), to examine degeneracy in functional neuroanatomy. Our model is built off of earlier topographic factor analysis approaches [21] and takes as input individuated segments of 4D fMRI timeseries data with labels for participant and trial. It does not require knowledge about the attributes of participants (demographic, personality, genetic, etc.), nor does it require knowledge about how trials sort into conditions. NTFA learns a low-dimensional representation - or an embedding - of functional activity for each participant and trial on the basis of shared patterns of neural activation from segments of data. The embeddings provide a simple, readily visualizable depiction of whether and how neural responses during a task vary across participants, trials, and participant by trial combinations.

In this paper, our goal is to validate NTFA using a simulation approach. Computational simulations are critical to test whether novel computational models are capable of performing as expected in principle, that is, under conditions with a known ground truth. In practice, the data generating mechanisms for functional neuroanatomy are rarely, if ever, known. That is why it is of particular importance in cognitive neuroscience to develop modeling approaches that are capable of providing insight as to whether there is likely to be degeneracy in functional neuroanatomy from the data alone and with minimal supervision. Using computational simulations, we first demonstrate the considerable shortcomings of applying the most commonly used “univariate” activation-based analytical approach in fMRI data analysis when there is degeneracy. In the typical form of this analysis, a general linear model is used to determine whether functional activity in a given voxel or brain region (i.e. set of voxels) is greater during trials from one experimental condition relative to a baseline condition. We then implement NTFA on simulated datasets with minimal assumptions about whether trials ought to be nested into particular task conditions, or participants into particular groups. Our deliverable is a demonstration of the ability of NTFA to recover embeddings that reveal degeneracy, and non-degeneracy, in simulated 4D timeseries data with topological structure (e.g. as in fMRI data).

## 2 Methods

Before discussing the model design in NTFA we begin by considering the consequences of applying widely used univariate analyses [22] to synthetic data that exhibit degeneracy. This approach serves two purposes. First, using synthetic data illustrates the pitfalls of using traditional univariate analyses in terms of capturing degeneracy. Second, this synthetic data provides a known ground truth to validate NTFA’s performance. Rather than extensively review the various forms of degeneracy that can occur in the brain, we decided to demonstrate two aspects of degeneracy that could occur in fMRI data. We generated the synthetic datasets to reflect a generic experimental framework in which participants undergo a baseline condition and an experimental condition. For example, in a study on fear, the baseline condition may consist of multiple trials that maintain a neutral affective state, and multiple trials that induce fear. In a study on working memory, there may be trials that involve low capacity demand in the baseline condition, and trials that involve high capacity demand in the experimental condition. We used the term, trial, to broadly represent trials in sequence (e.g. the first, second, …, trial of the task), or the specific contents of a trial in a task (e.g. trials that present stimulus A, stimulus B, …, in which each stimulus is a sampled instance from the same task). Degeneracy may occur in either case. For simplicity, each synthetic dataset consisted of two participants, and we assumed a single baseline state. We simulated multivariate patterns of neural activity throughout the brain by sampling from a prespecified underlying distribution. We then modeled three hypothetical situations to reflect different assumptions of degeneracy, which are described in more detail in the subsequent section(Figure 1). To demonstrate that standard neuroimaging analysis is limited in capturing degeneracy, we applied a standard univariate General Linear Model (GLM) to calculate a contrast between the experimental conditions and the baseline condition.

**Figure 1:**
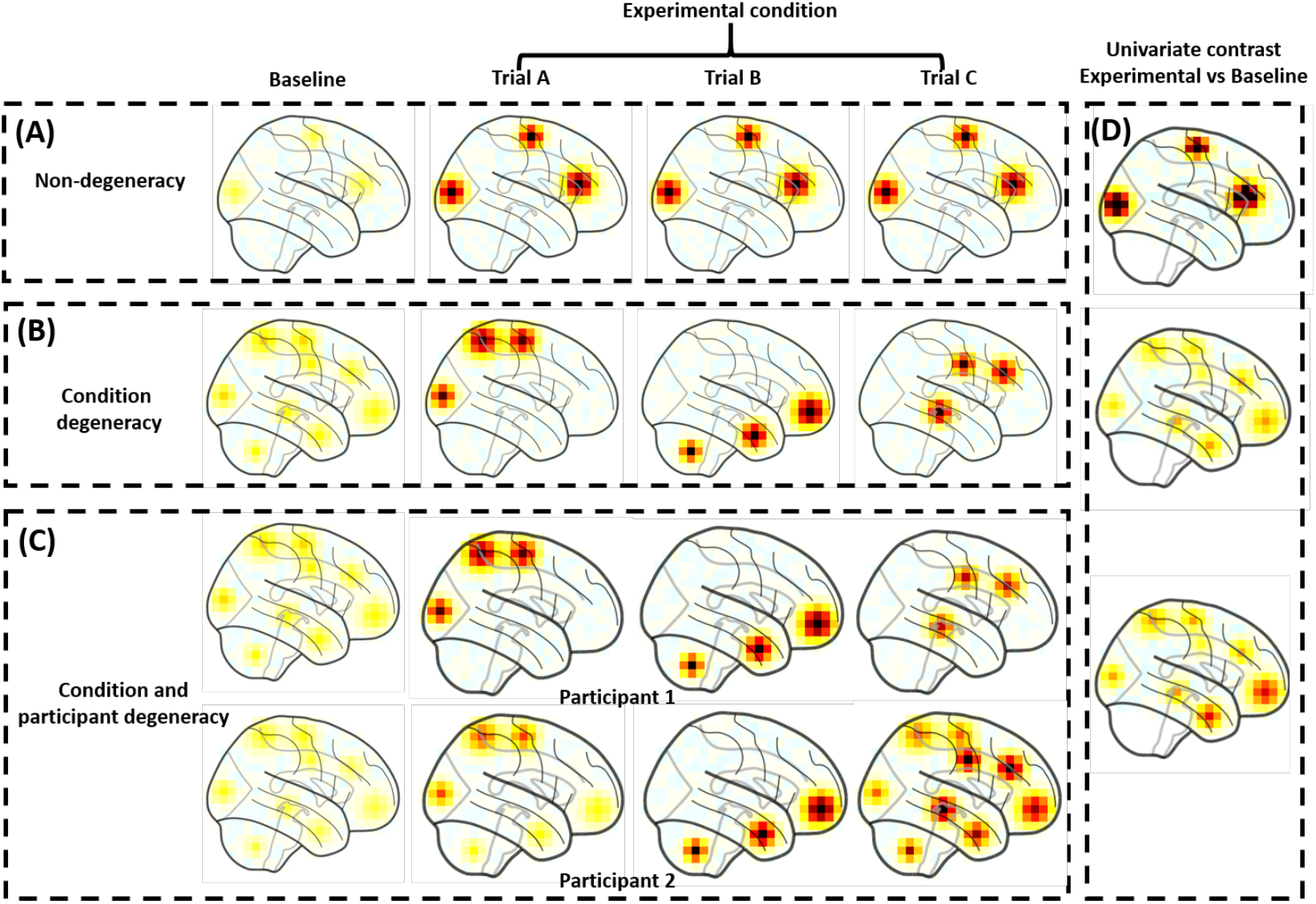
Standard univariate analysis applied to degenerate situations. We applied univariate analysis (right panel) to three simulated datasets (left panels), assuming a simple experimental design with a baseline condition and a task condition involving multiple trials. In an affective neuroscience task, for example, the experimental condition might be a fear condition, as designated and labeled by the experimenter, which consists of multiple trials that are thought to induce fear. **(A)** Non-degeneracy: We simulated data from a situation without degeneracy, in which a consistent set of regions are more active during the experimental condition than the baseline condition across trials (and across participants). **(B)** Condition degeneracy: Simulated data included different patterns of activation associated with different trials of the same experimental condition. **(C)** Degeneracy by condition and participant: Simulated data included different patterns of activation are associated with different trials and participants. **(D)** A traditional univariate analysis performs well in the situation without degeneracy. However, the analysis would be insensitive to the variations in the two situations involving degeneracy. Critically, with sufficient statistical power, the univariate analysis may still yield significant activations in situations B and C. However, the summary map would grossly mischaracterize the data, and the underlying data generating distribution.

### 2.1 Non-degeneracy

The non-degenerate functional neuroarchitecture stipulates that experimental trials evoking a common psychological state or process share a common underlying pattern of activation. We generated simulated data to fit this assumption. We started by selecting three brain areas randomly to create a pattern of activation during experimental condition trials (Figure 1A). We chose three areas arbitrarily to reflect the fact that the assumptions of a non-degenerate functional neuroanatomy have little to do whether the pattern of activation is localized to one area or distributed across many areas. What is important is that the same pattern of activation is assumed to occur consistently across trials and participants, and that a non-degenerate model treats variation as residual error. To capture this assumption in our synthetic data, we specified the data generating process as a unimodal distribution. This refers to one pattern of neural activity with some Gaussian distributed noise across trials and participants. The synthetic data from individual trials A, B, and C, as shown in (Figure 1A), were sampled from this distribution. This model suggests there is a common pattern of activation across all trials that evoke fear, for example.

We then evaluate how a standard univariate analysis using a GLM performs on this data. The GLM resembles a supervised analytical approach insofar as experimenters must specify beforehand the regressors in the model. In so doing, experimenters must make assumptions about how trials are nested into conditions. In our example experiment, Trials A, B, and C, would all be modeled with a single regressor since they belong to the same experimental condition. In that sense, the GLM shares the same assumptions as the data generating process. As shown in Figure 1D (top), applying the GLM to our synthetic data shows that it perfectly suits the non-degenerate functional neuroanatomy.

### 2.2 Degeneracy by Condition

Degeneracy by condition refers to the existence of multiple distinct patterns of neural activation that occur across trials of the same experimental condition. Using fear as our running example, different fear induction trials may involve different patterns of brain activation (Figure 1B). To simulate data corresponding to a degeneracy by condition model, our data generating process involved sampling from one of three different distributions. Each of the three distributions gave rise to distinct activation patterns from the others, while maintaining similar activation patterns within the distribution. In Figure 1B, Trials A, B, and C are exemplars, with each one sampled from a different distribution. Thus, degeneracy by condition suggests that multiple distinct activation patterns may occur during trials within the same experimental condition. There could be many reasons for degeneracy, as noted in the introduction and as we speculate upon in the discussion. However, the purpose of the synthetic dataset is to illustrate how well certain models would perform when the assumption of degeneracy by condition holds.

A standard univariate analysis does not perform well in this situation. Without knowledge of the actual data generating process, experimenters would again model the data using a single regressor for Trials A, B, and C – even though the underlying distributions are heterogeneous. In other words, the standard GLM requires the experimenter to make assumptions about how trials are organized into experimental conditions, with one of those assumptions being the absence of degeneracy. As a result, the GLM precludes the ability to test whether there is, or is not, a degenerate relationship. To illustrate the consequences, we fit a GLM to our simulated data. As shown in Figure 1D (middle), the univariate activation map appears as an amalgam of the three data generating distributions. Even when the ground truth (i.e. the underlying generative process) exhibits degeneracy by condition, a standard univariate analysis may still produce seemingly “reliable” findings (i.e. significant and reproducible findings with enough participants). However, the resulting pattern of activation in Figure 1D (middle) would not accurately capture the actual data generating process. Consequently, it could lead to a mistaken, but statistically “reliable”, conclusion about the relationship between neural activity and the experimental condition.

### 2.3 Degeneracy by Participant and Condition

For our third situation, we examined degeneracy with respect to both condition and participant. Similar to the example in the degeneracy by condition scenario, a participant would have different patterns of activation during different trials of the same experimental condition. In addition, however, the participant would also have a different pattern of neural activation than other participants, even during the same trial. For example, both participants may report experiencing the same level of fear when shown the same fear-inducing stimulus, but nevertheless show differential activation patterns.

This situation is illustrated in Figure 1C. Two participants may be presented with the same set of trial stimuli and even have the same behavioral responses, but the underlying neural patterns may nonetheless vary. For example, in Trial A, the exemplar data from two participants share activity in dorsal areas, but one participant also shows activity in ventral areas. In Trial B, they show similar patterns of activation. In contrast, in Trial C, there are again differences between participants. Thus, our data generating procedure was designed to capture: (i) degeneracy across participants by including both participant-specific activation patterns (e.g. Trials A and C), (ii) degeneracy by condition by including variation in activation patterns across Trials A-C within a participant, and also (iii) activation patterns that are also shared across participants (e.g. Trial B). We purposefully designed the synthetic data in this way to test the utility of NTFA in addressing this complexity.

Critically, degeneracy with respect to participants highlights another important assumption of standard univariate analyses. The analytical procedure of a GLM involves stages such that the outputs of the trial- and subject-level analyses are inputs to the group-level analyses. This sequence of analyses assumes a nested data structure in which trials of an experimental condition within one participant’s data and each participant from their group are from one normal distribution. This assumption is valid under a non-degeneracy functional neuroanatomy (shown in Figure 1A), but could preclude the ability to examine degeneracy in the functional neuroanatomy. Instead of applying the same first level model to all participants, a more appropriate model would fit the run and participant level simultaneously without assuming this nested structure.

Ultimately, the standard univariate approach are insensitive to variations across trials, compounded by degeneracy across individuals. It treats the systematic variation in activation patterns across trials and participants as error. Though it may produce reliable findings with sufficient power, it would result in a diffuse pattern of activation that is not representative of the data generating process. This is shown in Figure 1D (bottom) as applied to our synthetic data.

## 3 Neural Topographic Factor Analysis (NTFA)

In light of the shortcomings of the standard univariate analysis discussed in the previous section, there is a need for models that can uncover degeneracy when it is present in the data. We propose NTFA [23] for this purpose. NTFA is a generative model ^2^ built off of earlier topographic factor approaches for fMRI data [21] that is designed to learn low-dimensional, visualizable embeddings from segments of data for different participant-task combinations. The spatial positions of these embeddings can reveal different aspects of the data, including degeneracy. Moreover, NTFA is primarily unsupervised, requiring only the participant and trial identities ^3^. We provide an overview of NTFA’s generative model and training mechanism in Figures 2 and 3 respectively. A comprehensive explanation of these can be found in the supplement 5.

**Figure 2:**
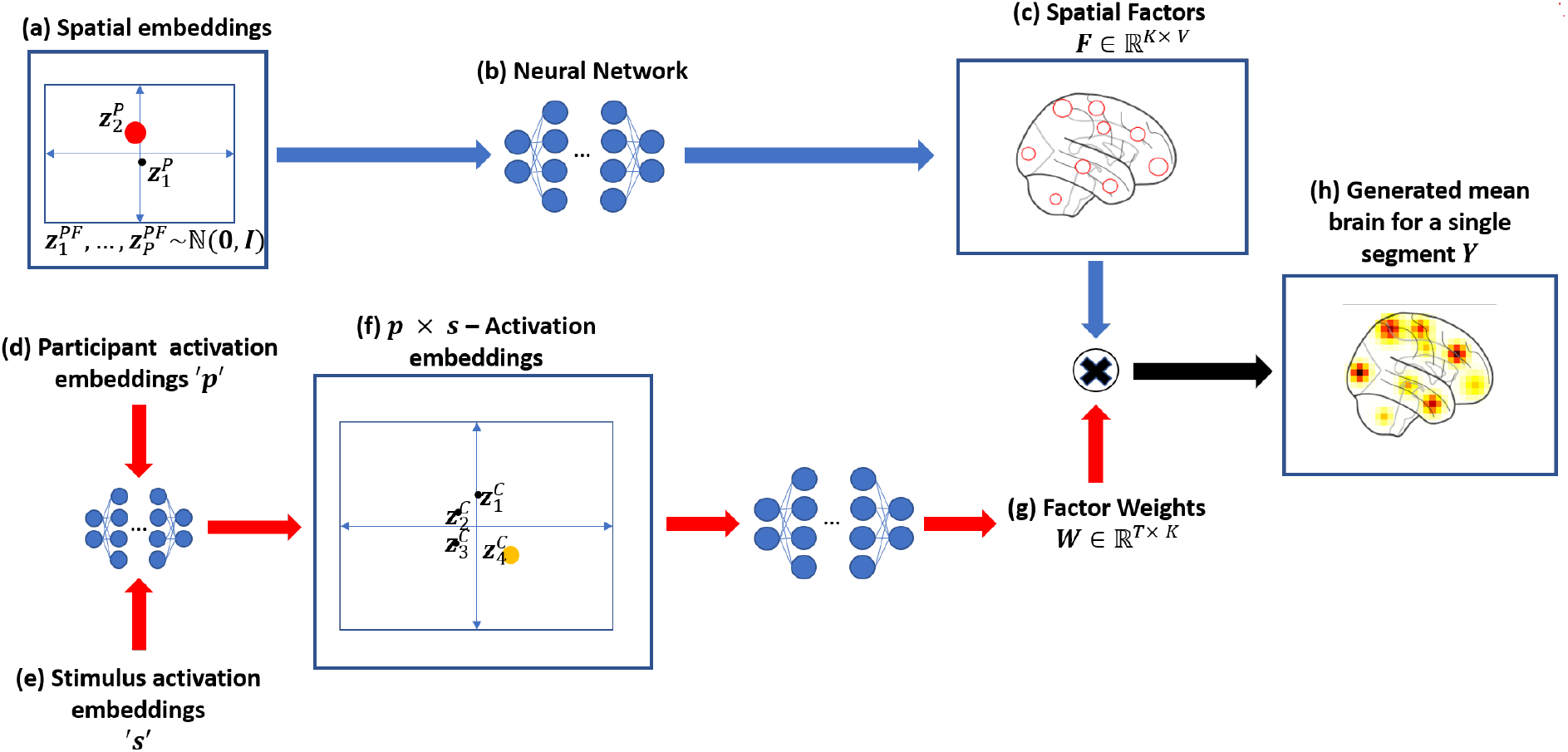
NTFA Generative Model: This figure describes how NTFA generates a single segment of fMRI data with *V* voxels and *T* TRs. NTFA treats a single participant-trial combination in the experiment as a segment of fMRI data such that it could model the participant and trial dependent activation without grouping participants or trials a priori. Concisely, NTFA splits this data generation into two parts, reflected by the two pathways in this figure. The first pathway, following the blue arrows, generates a participant dependent set of spatial factors. The second pathway, following the red arrows, generates the participant *and* trial dependent activation weights for these factors. The multiplication of these spatial factors and the factor weights gives us the generated fMRI segment. **(a-c) Generating spatial factors:(a)** We sample 2-dimensional spatial embeddings (*z*^PF^) from a gaussian prior, with each dot representing a participant in the shared embedding space. For each block we only use the spatial embedding for the participant in that block, shown here as the red dot. **(b)** This spatial embedding is submitted to a neural network. The same neural network is shared by all spatial embeddings. The use of neural networks allows a potentially non-linear mapping between the embedding space and the variations in the spatial factors.**(c)** The neural network maps this embedding to the *K* spatial factors to represent the functional units of activation in the brain, shown as the red circles. These spatial factors are assumed to be radial basis functions parameterized by the centers and widths output by the network. Here we show these spatial factors as red circles covering two widths of the radial basis function. The Spatial Factors is denoted by a matrix *F* of size *K* x *V*. As such, the differences in the spatial embeddings reflects the variations in these spatial factors. **(d-g) Generating factor weights:(d,e)** Similar to the spatial embeddings we also sample a participant activation embedding for the same participant and trial activation embedding for the trials across task conditions corresponding to the combination. These embeddings are meant to capture overall participant and trial dependent activity respectively. **(f)** These two embeddings are then passed to a neural network to produce the corresponding *p × s*- activation embedding. Each dot represents a unique participant and trial combination. **(g)** The activation embedding is then passed through another neural network to generate the Factor Weight matrix of *W* of size *T × K*. The factor weights capture the activations of the spatial factors. The neural network outputs the mean and a standard deviation of activation for each factor. Each factor’s activation is then generated by sampling independently over TRs from the corresponding Gaussian distribution to create the time varying weights *W*. As such, variations in locations of these activation embeddings reflects variations in the activations of spatial factors. The embeddings provide a way to visualize high dimensional variations between brain activations for different participant-stimulus combinations. **(h)** Finally, these weights and spatial factors can be arranged in the form of two matrices *W* ∈ ℝ^*T* ×*K*^ and *F* ∈ ℝ^*K*×*V*^. The matrix of spatial factors *F* and their activations *W* can be multiplied to generate data **Y** i.e. this segment of fMRI data. For a comprehensive version of this figure, see Figure. 5 in Appendix.

**Figure 3:**
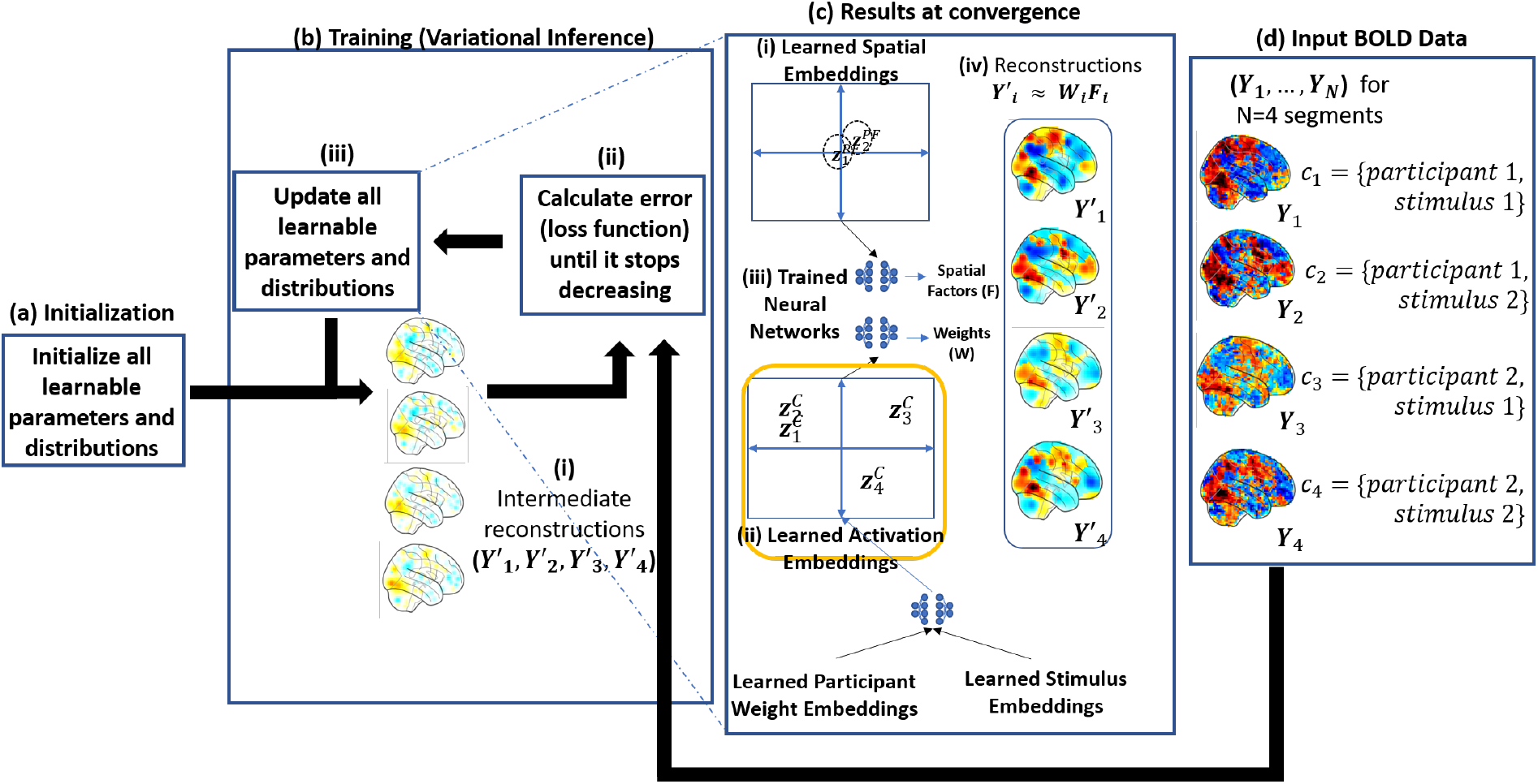
NTFA Training using variational inference: This figure shows the training procedure for NTFA for a hypothetical dataset that includes two participants and two stimuli for a total of four combinations. Mean brain images for the four segments can be seen in panel (d) where the preprocessed BOLD data is split into segments of participant-trial combinations, denoted here as *c*1*, c*2*, c*3 and *c*4 for this hypothetical example. **(a) Initialization** All parameters and distributions are initialized as specified in Appendix Section 5. **(b) Training (b-i)** All parameters and distributions are first initialized (see Appendix Section 5.). **(b-ii)** The parameters are used iteratively to calculate the reconstructions error. The loss function is defined as the sum of reconstruction error and the regularizer (see Equation 10 in Appendix Section 5). This is a consequence of using variational inference which aims to approximate the unknown posterior distributions of all the hidden variables with a set of simpler distributions, Gaussian in this case. **(b-iii)** These parameters are then updated in the direction of decreasing loss using stochastic gradient descent (SGD). The iterations are repeated until convergence that is when the loss function stops decreasing. **(c) Results at convergence** The learned parameters at convergence are represented by the embeddings. The embeddings provide a visual conclusion of variances in neural activity across different participant-trial combination. **(c-i)** The learned spatial embeddings encode the relative differences in the locations and widths of the spatial factors between participants. **(c-ii)** The learned activation embeddings are highlighted here in yellow as they are the main focus of this paper. These embeddings represent the differences in activation of the spatial factors among different participant-trial combinations. For example, in this hypothetical case the combinations 1 and 2 on the left of the plot are more similar to each other as compared to combinations 3 and 4. **(c-iii)** The three trained neural networks allow us to capture potentially nonlinear relationships between different participants’ spatial factors as well as activations for different combinations. These neural networks can also be used to generate unseen data including unseen participant-trial combinations by providing inputting appropriate embeddings. **(c-iv)** Shows the learned reconstructions that should approximate the major patterns in the input data as can be seen by side by side comparison with panel (d) with a limited number of spatial factors *K ≪ V*. For a comprehensive version of this figure see Figure 6 in the Appendix.

NTFA is designed to enable systematic comparison of functional neuroanatomy across individuals and task conditions by mapping fMRI data to low-dimensional (and visualizable) embeddings. We achieve this goal by formalizing three assumptions:

- First, we assume voxel-level data can be parsimoniously expressed as a much smaller set of functional units, which we refer to as **spatial factors**. We model these spatial factors as radial basis functions, and the activation at a given voxel as a sum of weighted contributions from these factors.
- Second, we assume that the same spatial factors exist in all participants, but their precise spatial location may vary across individuals. A set of low dimensional participant dependent **spatial embeddings**(*z*^PF^) capture this variation. A neural network maps these embeddings to the centers (location) and widths (extent) of the spatial factors. This neural network is shared across participants. The neural network allows us to learn a possibly nonlinear mapping from the space of spatial embeddings to that of spatial factors. This is important, as the anatomical alignment literature [24, 25] makes it implausible that this relationship can be captured with a linear transformation. Similarly, sharing a single neural network among all factors and all participants allows the spatial embeddings to be commensurable between participants. A Gaussian prior on the spatial embeddings encourages them to be close to each other.
- Third, we assume that degeneracy or non-degeneracy is effectively revealed as a combination of how a single participant’s brain responds to the various trials in a task condition (i.e. participant dependent activity) and how multiple individuals might respond to a the same trial in a task condition (i.e. trial dependent activity). By combining estimates of these sources of variation, we are able to detect whether neural activity in response to the same trial varies systematically across individuals, which we refer to as participant task combinations. Similar to the approach used for the spatial embeddings, participant dependent (*p*) and trial dependent (*s*) activity is estimated across the spatial factors (through embeddings *z*^PW^ and *z*^S^ respectively) and combined through a neural network to generate (**p × s**)**activation embeddings**(*z*^C^). The use of shared neural networks here once again ensures that the low dimensional embeddings can capture non-linear effects and make these embeddings commensurable between different participant-task combinations.

Taken together, these spatial and activation embeddings respectively provide a low-dimensional summary of where and how individuals’ brains respond to an experiment. Critically, the activation embeddings also summarize whether such responses are shared or diverge across individuals, hereby revealing potential degeneracy. For the simulated data from the three models discussed above, we can expect these activation embeddings to arrange in the following clearly different ways:

- **Non-degenerate**: For the non-degenerate scenario discussed in Section 2.1, we would expect the participant-trial activation embeddings to broadly fall in just two clusters: one for baseline and the other for the experimental condition. Figure 4(A) shows that the embeddings learned from NTFA indeed fall into two clusters.
- **Degeneracy by condition**: In the scenario discussed in Section 2.2, the activation embeddings are expected to fall in four distinct clusters: one for the baseline, and one each for the three underlying degeneracy modes. These will correspond to the differences in the three trials. Figure 4(B) shows that is indeed the case for the learned embeddings on this data.
- **Degeneracy by condition and participant**: In the scenario discussed in Section 2.3, the activation embeddings can be expected not only to group by trial, but also to split up by participants, with trials A and C revealing the degeneracy by condition and participants. Figure 4(C) shows precisely this expected behaviour.

**Figure 4:**
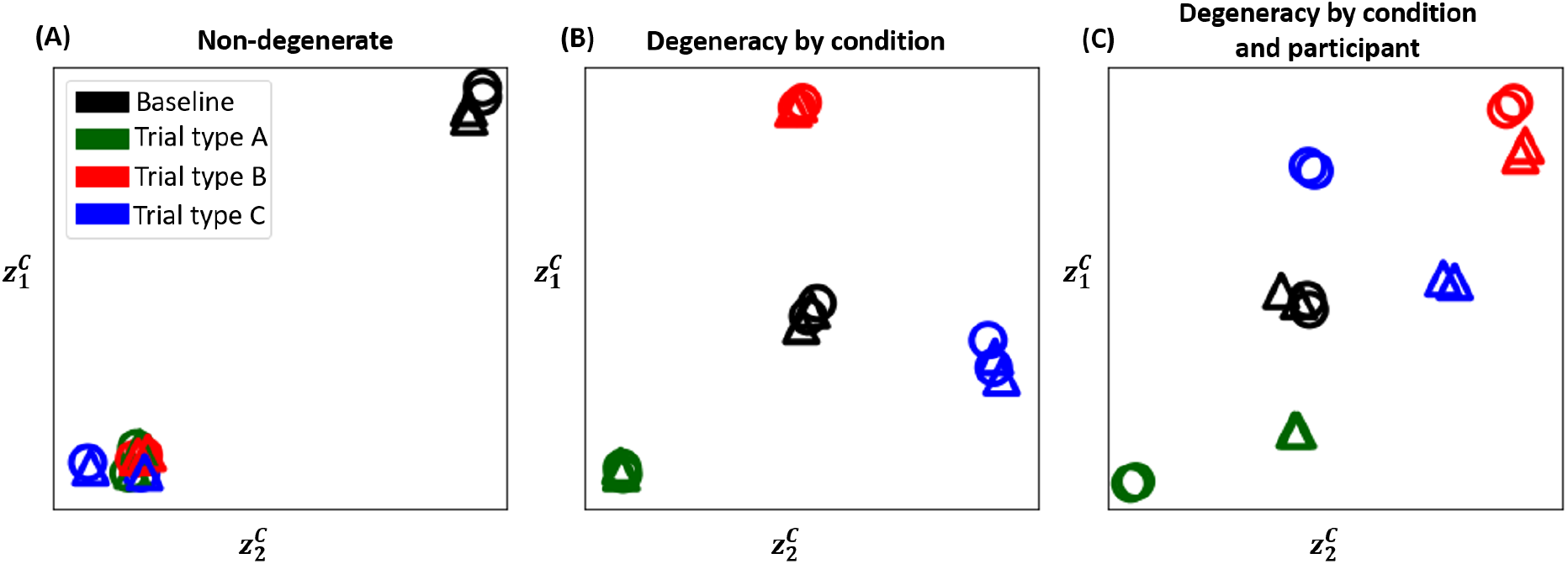
Inferred activation embeddings: The activation embeddings learned from NTFA for the three scenarios depicted in Figure 1 are shown here. NTFA was trained in an unsupervised manner and labels and colors are overlaid only for visualization and interpretation purposes. Each point represents a unique participant-trial combination. The colors correspond to trials as shown in the legend. Circles represent participant 1 and triangles represent participant 2. **(a) Non-degenerate:** The embeddings suggest there is no degeneracy, with combinations for all three experimental condition trials grouping together and away from the baseline combinations. **(b) Degeneracy by condition:** The embeddings suggest degeneracy in brain response based on trials, as the combination embeddings for each trial form a cluster of its own away from other clusters and away from baseline. There are no participant driven differences suggesting no degeneracy by participants. **(c) Degeneracy by condition and participants:** The embeddings here suggest degeneracy by both trials as well as participants, with the combinations forming groups of their own based on not just trials, but also splitting up by participants in case of Trial A and Trial C.

## 4 Discussion

Recent work in computational biology and functional neuroanatomy suggests that the brain may have multiple solutions, or degenerate neural pathways, when trying to solve a given task. However, current analytical methods are not optimized to capture such degeneracy. Here, we advanced a novel computational approach, NTFA, to address this issue. NTFA is a generative model that learns a low-dimensional space of embeddings from the temporal and spatial variation of fMRI data. The embeddings yield a visualizable representation of the latent variations in functional activity across trials and participants. The distribution of these embeddings can provide useful information for researchers to assess whether the data generating mechanism is degenerate or non-degenerate with respect to trial conditions and participants.

Related to NTFA, there are other models that also use latent factorization methods to analyze fMRI data, however, they are not currently equipped for modeling degeneracy with respect to task conditions and participants. These include topographic latent factorization models, such as topographic factor analysis and hierarchical topographic factor analysis [26, 21, 27], and non-topographic models such as principal component analysis [28], independent component analysis [29], the shared response model [30], hyper alignment [24], and dictionary learning methods [31]. These approaches are designed to address variation in functional alignment (e.g. individual differences in precise locations of the so-called fusiform face area [32, 25]). However, none of these methods explicitly model variation in functional activity across *both* participants and task conditions. Of note, NTFA is flexible in its implementation. If researchers preferred to label their trials as belonging to specific task conditions, or participants as belonging to specific groups, NTFA can accommodate these assumptions and develop a generative model with these assumptions built in (e.g. for more direct comparison with other approaches). NTFA’s other features may also be useful to the community. For example, NTFA explicitly models variation in the locations, sizes, and magnitudes of activation, whereas the vast majority of studies using univariate analysis of fMRI data focus only on activation magnitudes.

NTFA is, of course, not without some limitations, one of which is determining whether learned embeddings are modeling functionally meaningful signal or simply noise. Although much variation in fMRI data across time/trials (and across participants) is noise and should be discarded, that does not mean that all (or even most) variation unaccounted for by standard modeling approaches is necessarily noise. Here, we suggest that there is good reason to think that such variation might be structured and functionally meaningful (as described next), that historical approaches are insensitive to such variation unless it aligns with a narrow range of a priori hypotheses, and that NTFA is a technique that is designed to sift potentially interpretable, structured variation from random noise.

While our primary aim is constrained to establishing and validating our model using simulations, highlighting some relevant research findings may point to useful future directions in which to develop applications for NTFA. In general, it is well-known that psychological tasks are not “process pure” [33, 34]. A given task may involve a variety of different cognitive processes, neural pathways and/or strategies, which may shift and change over time and trials. Indeed, carefully constructed experiments have found results consistent with degeneracy even when using more traditional analytical tools. For example, dissociable neurocognitive memory systems can be used to complete the same overt memory task [35, 36, 37, 38]. When one system is compromised due to brain damage, other systems may be utilized to nonetheless complete the task at hand [39, 40, 12]. An increasing number of findings suggest that the brain is likely to offer multiple solutions in other domains too, such as in social cognition [41, 42] and emotion [43, 20]. NTFA may also be of particular relevance for translational research. Emerging work suggests that distinct neuropathologies may underlie a common clinical phenotype [44]. For example, research on depression suggests that there may be many different neuropathologies that give rise to depressive symptoms [45, 46, 47, 48]. Indeed, the call for “precision medicine" reflects a general failure of more traditional, non-degenerate theoretical models and rigid analytical approaches to account for heterogeneity in the underlying neural causes of mental health. A systematic evaluation of this variance is a critical step towards enabling precision medicine approaches in fMRI, in which neuroimaging studies have the potential to significantly advance diagnosis and treatment [49].

Despite these notable empirical examples, more often than not researchers assume that a given task involves a core set of processes that are shared across trials and participants. This may be because more traditional theoretical models in cognitive neuroscience rarely postulate degeneracy in functional neuroanatomy. However, more recent, predictive processing models of the brain suggest that degeneracy is likely to be common in mind-brain mapping [14, 50]. Another reason that researchers tend to assume a non-degenerate functional neuroanatomy is because it has been analytically challenging to not make this assumption. By addressing this analytical gap, NTFA offers new opportunities to model structured variance in fMRI data with a degree of independence from our own preconceived ideas of how this variance ought to be structured, and the opportunity to discover and model degeneracy in functional neuroanatomy.

## Acknowledgements

Research reported in this publication was supported by Department of Graduate Education (NCS 1835309) and the Brain and Cognitive Sciences Division (1947972) of the National Science Foundation.

## Author contributions statement

Z.K., Y.W., A.S. and J.B.H. designed the study. Z.K. and Y.W. performed the analysis and drafted the manuscript with guidance from A.S. and J.B.H. All authors reviewed and provided feedback on the manuscript.

## Footnotes

1 The concept of degeneracy may overlap with redundancy because they both suggest there are multiple solutions that can produce the same output, however they differ in the flexibility for the system to choose which solution to produce the outcome. See discussion in [1, 13, 17, 14].

2 A generative model here refers to a probabilistic model that characterizes a probability distribution from which new data could be generated in order to resemble the observed data

3 NTFA is flexible in the sense that users can incorporate supervision by specifying *a priori* groups of participants or trials. This can be done by providing the same identity to participants (or trials) belonging to the same group

