## Supplementary Materials for "A computational neural model for mapping degenerate neural architectures"

### 5 Details of Neural Topographic Factor Analysis

Neural Topographic Factor Analysis (NTFA) is a generative model for fMRI data. A generative model here refers to a probabilistic model that characterizes the probability distribution from which new data could be generated that resembles the observed data. Generative models are attractive primarily for three reasons: First, using generative models we can state our assumptions about the properties of the data in terms of hidden variables of interest. In this study, the observed data is the segments of an fMRI experiment and we are interested in the hidden structure of the data, i.e., how many different underlying groups of responses to trials are present in the data across participants. NTFA assumes that the data is generated by a non-linear mapping of low dimensional and visualizable embeddings that are participant and trial dependent. This non-linear mapping is assumed to be shared across all segments and therefore groups in the embedding space can be interpreted as the group structure present in the data and any variation from a group structure can also be revealed. We discuss NTFA's generative model in greater detail in Section 2.3.1 and Figure 2. The standard fMRI analyses like GLM on the other hand, often require the groups to be specified beforehand and then test for group level differences among them, individual variation is treated as error. This makes NTFA more suitable to investigate degeneracy in functional neuroanatomy as compared to traditional GLM like approaches that do not have a way to reveal degenerate responses that are different from the pre-specified groups.

Second, inference on generative models allows us to estimate the distributions of these hidden variables i.e. it provides uncertainties and not just point estimates. In case of NTFA, the hidden variables of interest include embeddings for participant and trial which generate embeddings for participant-trial combinations. Learning distributions of these embeddings rather than just point estimates means we can visualize the hidden structure in the data and specify our uncertainty about the learned structure. The ability to specify uncertainty is particularly useful in fMRI studies with limited data, as point estimates only become asymptotically correct as more and more data is available.

Lastly, learning a generative model allows us to generalize to unseen data. In our case, learning embeddings for each participant and trial and their generative mapping to the combination embedding space means we can predict these combination embeddings for participant-trial pairs that were not encountered during training. More broadly, this ability to make concrete predictions about unobserved data stands in contrast with traditional approaches in psychology that prioritize understanding explanatory mechanisms such as the GLM. The prediction-focused approach can generalize better to new data and is useful in many applied domains (Yarkoni & Westfall, 2017).

In the following sections, we first discuss NTFA generative model in detail. We then explain the variation distribution used for the inference procedure and how it is initialized. Lastly, we explain how the NTFA deploys the variational inference procedure iteratively to estimate the distribution of the latent variables given the observed data.

#### 5.1 Generative Model

The crux of NTFA's generative model can be explained in three parts. **First** is to assume that a block of fMRI data consisting of $T$ time points and $V$ voxels $Y \in \mathbb{R}^{T \times V}$ can be approximated by the matrix product of two matrices $Y \approx WF$; a matrix $F \in \mathbb{R}^{K \times V}$ that defines the spatial location of $K \ll V$ factors, with each row defining that factor's influence over each voxel, and a matrix $W \in \mathbb{R}^{T \times K}$ defining the weight of each factor at each time instant. **Second**, the model assumes that for a given participant "$p$" in a block, the parameters that define matrix $F$ can be generated from a lower dimensional vector $z_p^{\text{PF}}$ by passing it through a trainable non-linear mapping (a neural network $\theta_{\text{F}}$ in this case). This neural network is shared across all trials and all participants, which means the factors for all participants are generated through a shared mapping and the differences in the lower dimensional vectors can be interpreted as differences in matrix $F$ for participants across the experiment. **Third**, the model also assumes that given a participant-trial combination "$c = p \times s$" in a block, the parameters that generate the matrix $W$ for this block are generated by another lower dimensional vector $z_c^{\text{C}}$ mapped through another neural network $\theta_{\text{W}}$. This neural network is also shared across trials for all possible combinations and thus the differences in the lower dimensional vectors can be interpreted as differences in the activation of the spatial factors for different participant-trial combinations. This embedding is itself the output of a neural network $\theta_{\text{C}}$ that takes as input a participant dependent embedding $z^{\text{P}}$ and a task dependent embedding $z^{\text{S}}$. In the following paragraphs we unpack this model and the underlying assumptions in more detail. This description is also summarized and presented in Figure 5 for an example setting.

Let's assume we want to generate fMRI data for an experiment with $n = \{1, \ldots, N\}$ blocks. Each block $n$ consists of a participant $p_n$ out of a total of $P$ participants ($p_n \in \{1, \ldots, P\}$) undergoing a trial $s_n$ out of a total of $S$ unique trials ($s_n \in \{1, \ldots, S\}$). This leads to every block being defined by a combination $c_n = \{p_n, s_n\}$ of the participant identity and trial identity, where $c_n \in \{1, \ldots, C = PS\}$.

The first assumption we make is that each participant $p$ has a D-dimensional spatial embedding vector $z_p^{\text{PF}}$ (Figure 5A) and a participant embedding vector $z_p^{\text{P}}$ (Figure 5E) associated with it. The participant embedding is the vectors of all participants plotted in a 2-dimensional space. The spatial embedding captures the mean and variance of the center and width for each spatial factor in the brain space. The participant embeddings captures the participant dependent response across all trials in a task condition, Similarly we assume that each trial $s$ also has a separate D-dimensional trial embedding vectors $z_s^{\text{S}}$ Figure 5F) associated with it. We assume $D = 2$ for both cases as we would like to be able to visualize these vectors. These embeddings allow us to reason about differences between participants and trials as signal rather than noise. These participant and trial embeddings then pass through a neural network $\theta_{\text{C}}$ to generate participant-trial activation embeddings $z^{\text{C}}$. These combination embeddings in turn generate through another neural netwok $\theta_W$ the parameters for the distributions of activations of the spatial factor for a given participant-trial combination.

The second assumption is that these embeddings are sampled from a standard normal prior (a gaussian distribution with zero mean and identity covariance i.e. $\mathcal{N}(0, I)$). The embeddings are assumed to lie in two *separate* 2-dimensional spaces as shown in Figure 5(A, E, F). Note that we will infer the distributions of each of these embeddings later, these priors serve to constrain the space in which these embeddings lie in relation to each other.

$$z_p^{\text{P}} \sim \mathcal{N}(0, I), \qquad z_p^{\text{PF}} \sim \mathcal{N}(0, I), \qquad z_s^{\text{S}} \sim \mathcal{N}(0, I). \tag{1}$$

The third assumption is that the participant weight embeddings $z_p^{\text{P}}$ and trial embeddings $z_s^{\text{S}}$ can be combined through a non-linear mapping (with a simple neural network) to generate the combination embedding $z_c^{\text{C}}$ for that particular participant-trial combination.

$$z_c^{\text{C}} \leftarrow \theta_{\text{C}}(z_p^{\text{P}}, z_s^{\text{S}}), \tag{2}$$

The fourth and the most critical assumption is that the spatial embeddings and the activation embeddings can be mapped to two matrices: a matrix of factors $F \in \mathbb{R}^{K \times V}$ and a matrix of weights $W \in \mathbb{R}^{T \times K}$ through a non-linear mapping (using neural networks). Where $V$ is the number of voxels in the fMRI data and $T$ is the number of time points in a block. To realize this mapping, we assume that after sampling a participant embedding $z_p^{\text{PF}}$ using Equation (1) it can be passed through a neural network $\theta_{\text{F}}$ that outputs four quantities. It outputs 3-dimensional means of $K$ centers $\mu_p^x$ in voxel space, 3-dimensional standard deviations $\sigma_p^x$ associated with these means. Similarly it outputs 1-dimensional means of $K$ log-widths $\mu_p^\rho$, and associated 1-dimensional standard deviations $\sigma_p^\rho$ (Figure 5(c)). After generating these means and standard deviations, we assume that the $K$ centers for the participant $p$ i.e. $x_p^{\text{F}}$ and $K$ log-widths $\rho_p^{\text{F}}$ can be sampled from Gaussian distributions with means and variances generated above (Figure 5(d)).

$$x_p^{\text{F}} \sim \mathcal{N}(\mu_p^x, \sigma_p^x), \qquad \mu_p^x, \sigma_p^x \leftarrow \theta_{\text{F}}(z_p^{\text{P}}), \tag{3}$$

$$\rho_p^{\text{F}} \sim \mathcal{N}(\mu_p^\rho, \sigma_p^\rho), \qquad \mu_p^\rho, \sigma_p^\rho \leftarrow \theta_{\text{F}}(z_p^{\text{P}}). \tag{4}$$

Once the centers and log-widths are sampled using Equation (3) we can use these to define $K$ spatial factors using a radial basis function. That is, each factor $f_k$ is defined as a Gaussian "blob" centered at $x_{p,k}^{\text{F}}$ with a log-width $\rho_{p,k}^{\text{F}}$. Each factor $f_k$ defines a single $V$-dimensional row of the matrix $F_p$ for participant $p$ (Figure 5(e)).

Note that the neural network $\theta_{\text{F}}$ is the same for all participants, implying that this mapping is shared across participants and for all blocks. The embedding $z_p^{\text{PF}}$ once sampled for a particular participant also stays

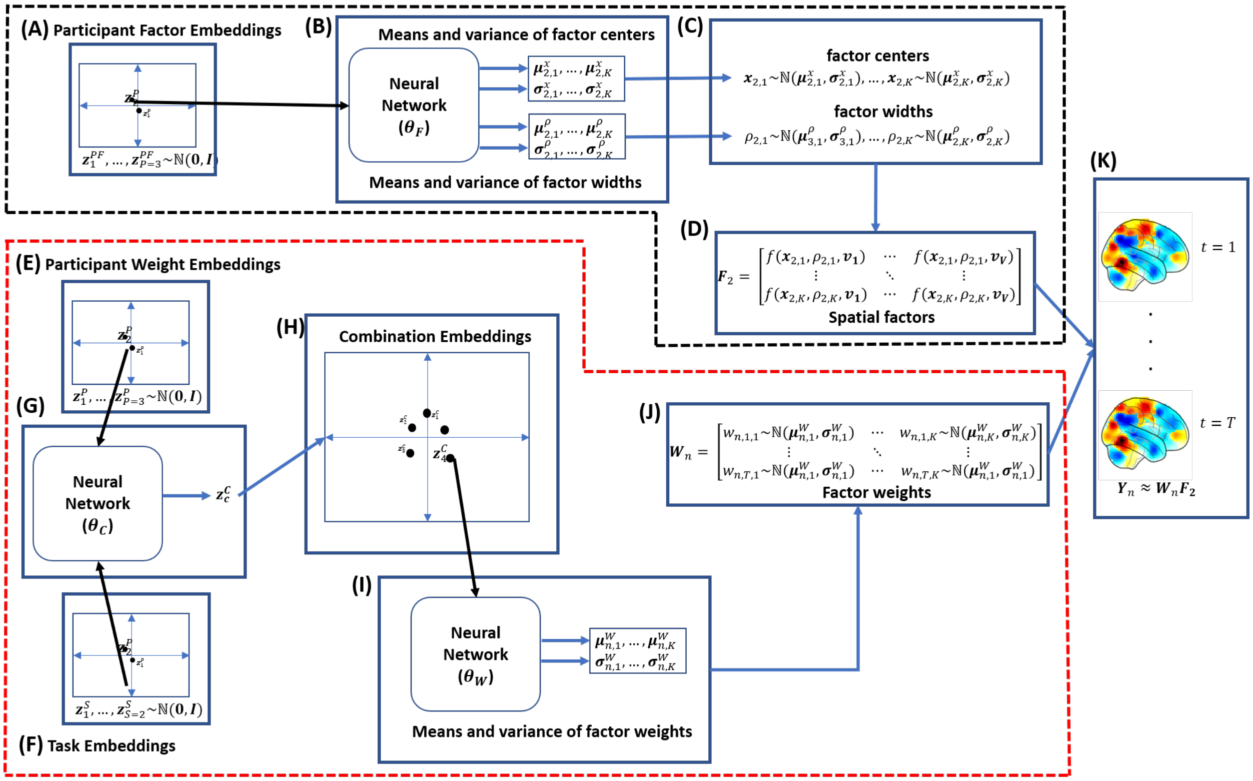


Figure 5: **Generative Model of NTFA.** The figure illustrates NTFA's generative model. For ease of visualization we consider the case with $P = 3$ participants, $S = 2$ unique trials and therefore $C = 6$ unique participant-trial combinations. The generative model can be divided into two main parts. The first part outlined in black dashed lines generates the spatial factors ($F$). The second part (inside red dashed outline) generates the weights ($W$) of these factors. Matrix product of $W$ and $F$ gives us the data $Y$. **(A)** The participant factor embeddings $z_1^{\mathrm{PF}}, z_2^{\mathrm{PF}}, z_3^{\mathrm{PF}}$ are sampled from a standard Gaussian prior. **(B)** The neural network $\theta_{\mathrm{F}}$ takes that participant's embedding $\mathbf{z}_2^{\mathrm{P}}\mathrm{F}$ as an input and outputs a set of parameters; the means and variances of centers $\mathbf{x}$ and widths $\rho$ for each of $K$ spatial factors. **(C)** For each factor, the centers and widths are then sampled from a gaussian distribution with mean and variances from the output of the neural network $\theta_{\mathrm{F}}$. **(D)** The spatial factors for this participant are now constructed from these sampled factor centers and factor widths by using radial basis functions $rbf(x, \rho, v)$. Each row of the factor matrix $F_3$ describes one spatial factor. The $rbf$ function takes as input the center of a factor $x$, the width $\rho$ and a voxel location $v$ to calculate the spatial coverage of the factor at voxel location $v$. **(E)** Participant weight embeddings $z_1^{\mathrm{P}}, z_2^{\mathrm{P}}, z_3^{\mathrm{P}}$ are also sampled from a standard Gaussian prior. **(F)** Similarly, trial embeddings $z_1^{\mathrm{S}}, z_2^{\mathrm{S}}$ are also sampled from a standard Gaussian prior. **(G)** The neural network $\theta_{\mathrm{C}}$ takes as input a trial embedding and a participant embedding sampled previously and generates the corresponding combination embedding. **(H)** Repeating the process in part (G) for all possible participant-trial pairs results in the combination embeddings shown here. **(I)** The neural network $\theta_{\mathrm{W}}$ takes the appropriate embedding $z_4^{\mathrm{C}}$ as input and outputs the means and variances for the weights of the spatial factors. **(J)** The factor weights at each time instant $t = 1, \ldots, T$ are then calculated by sampling from a gaussian distribution with the appropriate mean and variance for that factor. **(K)** The fMRI data $Y_n$ is then assumed to be approximately a matrix product of $W_n$ and $F_3$ with additive gaussian noise $\sigma^Y$. This same process can be repeated for any block for other participants $p$ and combinations $c$ to create a whole dataset with $N$ blocks.

the same across all blocks. These two assumptions combined indicate that there's something common for a participant across the whole experiment, and that the embeddings for the participants can be compared with each other.

Similarly we assume after generating the activation embeddings $z_c^{\mathrm{C}}$ using Equation (1) for a trial $n$ these can be passed through another neural network $\theta_{\mathrm{W}}$ to generate 1-dimensional means of $K$ factor weights $\mu_n^{\mathrm{W}}$ and associated standard deviations $\sigma_n^{\mathrm{W}}$. Then the weight for each factor can be sampled from a Gaussian distribution with the generated mean and standard deviation for each time point $t$ (Figure 5(f),(g)).

$$W_{n,t} \sim \mathcal{N}\left(\mu_n^{\mathrm{W}}, \sigma_n^{\mathrm{W}}\right), \qquad \mu_n^{\mathrm{W}}, \sigma_n^{\mathrm{W}} \leftarrow \theta_{\mathrm{W}}\left(z_c^{\mathrm{C}}\right). \tag{5}$$

Once we have $W_{n,t}$ and $F_p$ for a block our last assumption is that noisily sampling the matrix product of these two matrices generates the fMRI image at time $t$ for block $n$ (Figure 5(h)).

$$Y_{n,t} \sim \mathcal{N}\big(W_{n,t} F_p, \sigma^{\mathrm{Y}}\big), \qquad F_p \leftarrow \mathrm{RBF}(x_p^{\mathrm{F}}, \rho_p^{\mathrm{F}}). \tag{6}$$

This generative model can be summarized in the form a joint probability density over all the random variables in the model $p_\theta(Y, W, x^{\mathrm{F}}, \rho^{\mathrm{F}}, z^{\mathrm{P}}, z^{\mathrm{PF}}, z^{\mathrm{S}})$ which can be defined as follows:

$$p_\theta(Y, W, x^{\mathrm{F}}, \rho^{\mathrm{F}}, z^{\mathrm{P}}, z^{\mathrm{PF}}, z^{\mathrm{S}}) = p(Y \mid W, x^{\mathrm{F}}, \rho^{\mathrm{F}}) p_{\theta_{\mathrm{W}}}(W \mid z^{\mathrm{C}} = \theta_{\mathrm{C}}(z^{\mathrm{P}}, z^{\mathrm{S}})) p_{\theta_{\mathrm{F}}}(x^{\mathrm{F}}, \rho^{\mathrm{F}} \mid z^{\mathrm{PF}}) p(z^{\mathrm{S}}) p(z^{\mathrm{P}}) p(z^{\mathrm{PF}}) \tag{7}$$

### 5.2 Inference

The generative model we have discussed so far and summarized in Equation 7 describes the generation of the data. While the actual quantity of interest for us is what we can learn when we already have the data. Given data $Y$ from an fMRI experiment, all the other random variables in (7) are unobserved (latent) and we'd like to learn the distribution of these latent variables given the data i.e. we are interested in the posterior distribution $p_\theta(W, x^{\mathrm{F}}, \rho^{\mathrm{F}}, z^{\mathrm{P}}, z^{\mathrm{PF}}, z^{\mathrm{S}} \mid Y)$. Unfortunately, learning this distribution directly is intractable since it involves multiple integrations over all possible values of all the latent variables (See: Supplementary Information Bayes Rule). Fortunately, there is a group of techniques in Machine Learning literature called Variational Inference that aim to approximate the posterior distribution with a simpler distribution defined over all the latent variables. This approximate posterior distribution is often called variational distribution and denoted as $q_\lambda$ with parameters $\lambda$.

This variational distribution is often assumed to be factorizable, in our case this means assuming a variational distribution that is the product of individual distributions defined over all the latent variables as follows:

$$q_\lambda(W, \rho^{\mathrm{F}}, x^{\mathrm{F}}, z^{\mathrm{P}}, z^{\mathrm{PF}}, z^{\mathrm{S}}) = \prod_{n=1}^{N} \prod_{t=1}^{T} q_{\lambda_{n,t}^{\mathrm{W}}}(W_{n,t}) \prod_{s=1}^{S} q_{\lambda_s^{\mathrm{S}}}(z_s^{\mathrm{S}}) \prod_{p=1}^{P} q_{\lambda^{\mathrm{x}_p^{\mathrm{F}}}}(x_p^{\mathrm{F}})\, q_{\lambda^{\rho_p^{\mathrm{F}}}}(\rho_p^{\mathrm{F}})\, q_{\lambda_p^{\mathrm{P}}}(z_p) q_{\lambda_p^{\mathrm{PF}}}(z_p^{\mathrm{PF}}). \tag{8}$$

where $q_{\lambda_{n,t}^{\mathrm{W}}}(W_{n,t})$ approximates the posterior distribution of factor weights for trial $n$ and time point $t$. $q_{\lambda_s^{\mathrm{S}}}(z_s^{\mathrm{S}})$ approximates the posterior distribution of trial embedding for trial $s$. $q_{\lambda^{\mathrm{x}_p^{\mathrm{F}}}}(x_p^{\mathrm{F}})$ approximates the posterior distribution of factor centers for participant $p$, while $q_{\lambda^{\rho_p^{\mathrm{F}}}}(\rho_p^{\mathrm{F}})$ does the same for factor log-widths. $q_{\lambda_p^{\mathrm{P}}}(z_p)$ approximates the posterior distribution for the participant embedding for participant $p$ and $q_{\lambda_p^{\mathrm{PF}}}(z_p^{\mathrm{PF}})$ does the same for participant facto embedding .

Once we have defined the variational distribution in Equation (8) the next step is to learn the parameters $\lambda = \{\lambda^{\mathrm{W}}, \lambda^{\mathrm{S}}, \lambda^{\mathrm{X}}, \lambda^{\rho}, \lambda^{\mathrm{P}}, \lambda^{\mathrm{PF}}\}$ of this distribution and the neural network parameters $\theta = \theta_{\mathrm{W}}, \theta_{\mathrm{F}}, \theta_{\mathrm{C}}$ such that it comes as close as possible to the true posterior $p_\theta(W, x^{\mathrm{F}}, \rho^{\mathrm{F}}, z^{\mathrm{P}}, z^{\mathrm{PF}}, z^{\mathrm{S}} \mid Y)$. Once again using well known derivations (detailed in supplementary materials) this can be done without knowing the actual posterior distribution by instead maximizing the following objective with respect to $\lambda$ and $\theta$:

$$\mathcal{L}(\theta, \lambda) = \mathbb{E}_q \left[ \log \frac{p_\theta(Y, W, x^{\mathrm{F}}, \rho^{\mathrm{F}}, z^{\mathrm{P}}, z^{\mathrm{PF}}, z^{\mathrm{S}})}{q_\lambda(W, x^{\mathrm{F}}, \rho^{\mathrm{F}}, z^{\mathrm{P}}, z^{\mathrm{PF}}, z^{\mathrm{S}})} \right] \tag{9}$$

The right hand side of this equation can be split into two parts:

$$\mathcal{L}(\theta, \lambda) = \underbrace{\mathbb{E}_q[\log p(Y|W, x^{\mathrm{F}}, \rho^{\mathrm{F}})]}_{\text{negative of reconstruction error}} - KL(q_\lambda(W, x^{\mathrm{F}}, \rho^{\mathrm{F}}, z^{\mathrm{P}}, z^{\mathrm{PF}}, z^{\mathrm{S}}) \mid\mid p_\theta(W, x^{\mathrm{F}}, \rho^{\mathrm{F}}, z^{\mathrm{P}}, z^{\mathrm{PF}}, z^{\mathrm{S}})) \tag{10}$$

Since $p(Y|W, x^{\text{F}}, \rho^{\text{F}})$ is a Gaussian distribution, the first time on the right is equivalent to the negative of the expected reconstruction error between the observed data and the data reconstructed from the samples from the variational distribution $q_\lambda$. The second term is a regularizer term that measures how similar the variational distribution is to the prior distribution. Maximizing this objective with respect to $\lambda, \theta$ then equates to minimizing the reconstruction error as well as making sure that the priors and the variational distribution become similar.

This objective can be optimized using black-box methods provided by available libraries such as Probabilistic Torch [51]. Broadly this optimization proceeds in two steps, the first is to initialize the parameters of the variational distribution $q_\lambda$ and second is to sample from this $q$, calculate the objective (10) and then to iteratively update all parameters of $q$ in such a way that the objective is expected to increase until it stops increasing. We now discuss these two steps in the following paragraphs:

### 5.3 Initializing Variational Distribution

All distributions are assumed to be gaussian, owing to the universality of gaussian distributions and the ease of sampling and optimizing objective (10) when using gaussian distributions. This is also a fairly established standard practice in variational inference. Below we provide a list of how the means and variances of these gaussian distributions are initialized.

- The variational distributions over the participant embeddings $q_{\lambda^{\text{P}}_p}(z_p)$ $q_{\lambda^{\text{PF}}_p}(z_p)$ for a participant $p$ and trial embeddings $q_{\lambda^{\text{S}}_s}(z_s)$ for a trial $c$ are both initialized with a zero mean and unit variance. i.e. a standard normal. When we learn these distributions, we will not only learn a point estimate for these embeddings, but also an estimate of our uncertainty about the location of each embedding. The same initialization is used for all participants, and all combinations.

- The means of variational distribution over the centers of the factors $q_{\lambda^{x^{\text{F}}_p}}(x^{\text{F}}_p)$ and the means of variational distribution over factor log-widths $q_{\lambda^{\rho^{\text{F}}_p}}(\rho^{\text{F}}_p)$ can be initialized in two ways suggested by [21]: **1.** The means of centers can be initialized by performing k-Means clustering on the voxel locations using number of factors $K$ as number of clusters. The centers of the resulting clusters can then be used to initialize the means of factor centers. Each voxel is then labeled by the center closest to it and the variance of each cluster is used to initialize the mean of the width of each factor. **2.** By hotspot initialization, While this process is described in more detail in the supplementary material it involves placing the initial factor centers one by one at the peak of average fMRI image calculated from the whole dataset, solving a least square problem to approximate the width of that factor, subtracting this factor from the mean image and choosing the next peak as the next center until all factors have been initialized. The hotspot initialization works well for smaller number of factors for example when dealing with simulated data. A standard deviation of 1 is used to initialize the standard deviation of the variational distributions for factor centers. For factor widths, the standard deviation is initialized as the standard deviation of widths for all factors.

- The means of variational distribution for weights $q_{\lambda^{\text{W}}_{n,t}}(W_{n,t})$ are initialized by constructing the initial spatial factors using the centers and log-widths from the previous step (using a radial basis function), and then solving an ordinary least squares (OLS) problem. The OLS problem uses the average brain image computed across the whole dataset, and tries to learn the weights of the initial factors such that the weights and the factors combine can approximate this average image. The resulting weights are then used as mean of variational distributions for weights for all blocks $n$ and time points $t$. Once again the standard deviation is initialized to 1.

### 5.4 Training

Once the variational distribution $q_\lambda$ has been initialized, we can sample from this distribution and approximate the objective (10). At first iteration we sample the variables $W, x^{\text{F}}, \rho^{\text{F}}, z^{\text{P}}, z^{\text{PF}}, z^{\text{S}}$ from the initialized distributions for factor weights, factor centers, factor log-widths, participant embeddings and combination embeddings. This and the initial (random) weights of the neural networks are used to calculate the objective (10). This is equivalent to calculating the reconstruction error between the input data and the data

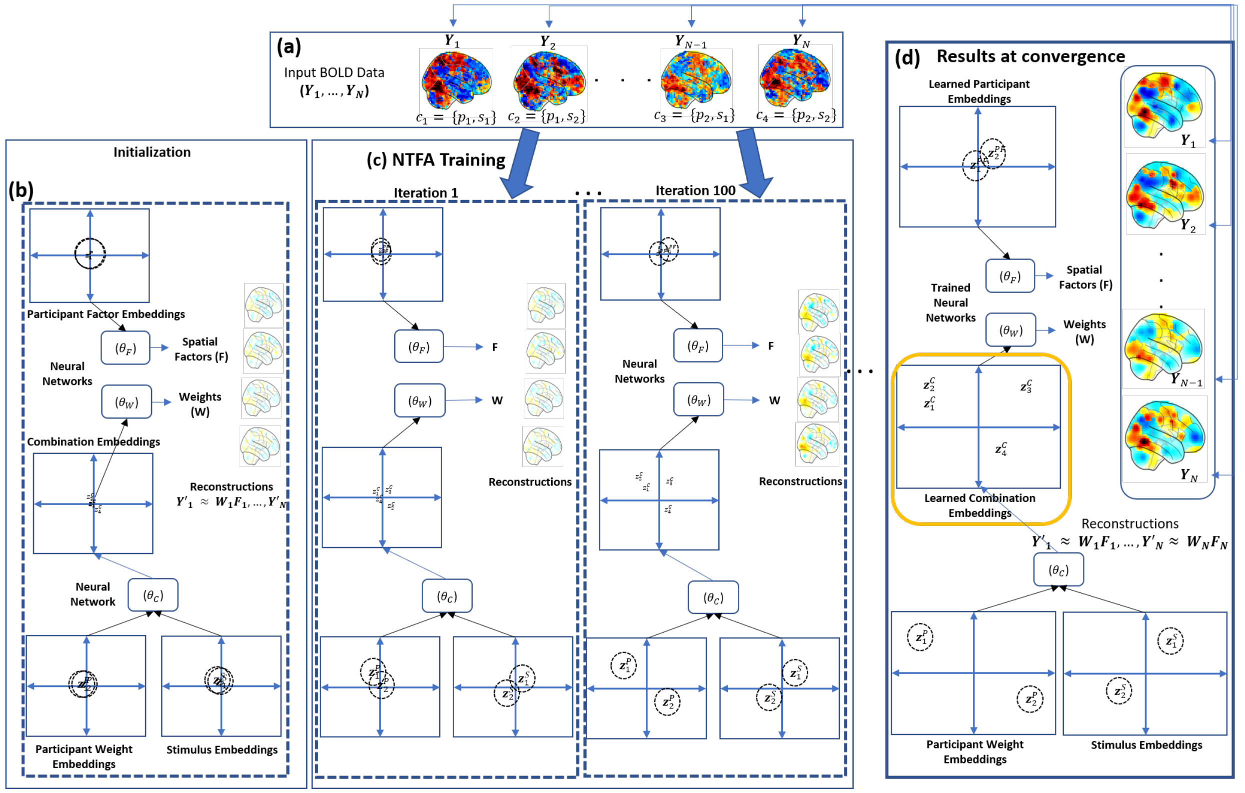


Figure 6: **NTFA Training.** **(a)** During training NTFA takes as input an fMRI dataset with $N$ trials, $Y_1, \ldots, Y_N$, for illustration purposes this dataset is assumed to have $P = 2$ participants and $S = 2$ unique trials, which implies $C = 4$ unique participant-trial combinations, note that a combination can occur multiple times in an experiment. **(b)** Variational distributions for all the latent variables are initialized as explained in Section 5.3. **(c)** The model trains by iteratively optimizing the variational distributions for the participant factor embeddings ($z^{\mathrm{P}}$), participant weight embeddings (($z^{\mathrm{PF}}$)) and trial embeddings ($z^{\mathrm{S}}$) as well as parameters of spatial factors and weights, and the three neural networks ($\theta_{\mathrm{F}}, \theta_{\mathrm{W}}, \theta_{\mathrm{C}}$). $\theta_{\mathrm{C}}$ takes as input the participant weight embeddings and the trial embeddings to generate combination embeddings $z^{\mathrm{C}}$. $\theta_{\mathrm{F}}$ and $\theta_{\mathrm{W}}$ generate the parameters of spatial factors and weights such that at each iteration the model is better able to reconstruct the original fMRI data. We show here the first iteration and $100^{th}$ iteration. As $100^{th}$ iteration the model is starting to learn the relative locations of the embeddings and the reconstructions are looking sharper as compared to iteration 1. **(d)** Once the training converges, the latent variables stop changing. The participant factor embeddings are still close to each other with overlap, suggesting the participants don't differ significantly in terms of the spatial factors. The participant weight embeddings are in clearly different regions of the space with no overlap, suggesting significant differences in the factor weights for the two participants. Similarly, the trial embeddings also suggest a significant difference between the two trials in terms of evoked response. The combination embeddings (highlighted in yellow) capture the relative differences in participants' brain response to each task. In this case, they suggest the first two combinations $c_1, c_2$ are more similar to each other as compared to $c_3, c_4$. $c_3$ and $c_4$ also appear to be different from each other. Lastly, the reconstructions at this point are reasonable approximations of the input data.

reconstructed from the the sampled factor weights and spatial factors, and a regularizer term that calculates the KL divergence between model prior distribution and the variational distribution. The parameters of the variational distribution $\lambda$ and the parameters of the neural networks $\theta$ are then updated using stochastic gradient descent in a direction that improves the expected reconstruction error in the next iteration and also makes the model priors and the variational distribution more similar. This process ensures that variational distribution is updated in such a way that samples from it can reconstruct the data well, at the same time the neural network parameters are updated in such a way that samples generated from the model will be more and more similar to the samples from the variational distribution. This process is repeated until convergence which is achieved when the value of the objective function in equation (10) stops changing for successive iterations. Once convergence is achieved we can analyze the posterior distributions of the participant embeddings and the combination embeddings by visualizing their means and standard deviation as shown in Figure **??**. We can also visualize the reconstructions by combining the posterior estimates of weights and factors. Similarly, at this point the neural network $\theta_{\mathrm{C}}$ is trained to generate combinations which in turn can generate average reconstructions for a block through the trained neural networks $\theta_{\mathrm{W}}, \theta_{\mathrm{F}}$ and can also be used to generate data similar to the training data by providing embeddings as input.

### 5.5 General Linear Model

To analyze the data, we implemented the GLM in which each trial was modeled as a separate regressor, such that the model estimated a statistical map for each trial. The model then calculated a contrast on trials in the experimental condition and on trials in the baseline condition to assess which voxels showed greater activity in the experiment conditions than in the baseline condition. We ran a one-sample t-test at every voxel to test which voxels' coefficients significantly differed from zero. The statistical map of the t-test over two participants was presented. The model did not include nuisance regressors and were not convolved with a hemodynamic response function.
